# B-REAL: A keypoint-less behavior platform for offline classification and real-time detection

**DOI:** 10.64898/2026.10.13.758637

**Authors:** A Eagle, B Kohler, V Kamrowski, N Lingam, R Li, PG Middlebrooks, EA Yttri

## Abstract

Two obstacles limit automated behavior analysis in neuroethology: the burden of first training pose estimation models, and the inability to identify behavior quickly enough for real-time classification that could trigger stimuli, drug delivery, or environmental manipulations. Here we present B-REAL, a pose-free classifier that maps raw video to behavior faster than video frames are recorded, even on a CPU. Given substantially less training data, B-REAL matched or surpassed established methods trained on full benchmark rodent or fly datasets, particularly on rare and even single-frame behaviors. Pretraining on unlabeled video further improves its accuracy without slowing inference or training time. In closed loop, rewarding mice for B-REAL-detected grooming more than doubled their performance of grooming bouts. B-REAL thus makes closed-loop behavioral experiments practical with minimal annotation and standard hardware.

## INTRODUCTION

Behavior is the primary observable output of nervous system function, and its precise measurement is essential for linking neural circuits, genetic variation, and pharmacological interventions to functional outcomes (Anderson and Perona, 2014; Datta et al., 2019). Historically, this quantification has relied on trained human observers scoring video recordings frame-by-frame — a methodology that, while flexible and interpretable, is inherently time-consuming and subject to inter-rater variability (Luxem et al., 2023; Goodwin et al., 2024). These limitations have constrained behavioral science to brief observational windows and coarse temporal resolutions, foreclosing entire classes of questions, including real-time integration of behavioral state with neural recordings or perturbations, a central and powerful theme in behavioral neuroscience especially in recent years (Yttri and Dudman, 2016; Zhang et al., 2025).

The past decade has seen remarkable progress in the automation of animal behavior analysis, driven largely by advances in deep learning and computer vision (Berman et al., 2014; Luxem et al., 2022; Hu et al., 2023; Weinreb et al., 2024; Choudhary et al., 2026; Daruwalla et al., 2026; Fong et al., 2026). The dominant paradigm that has emerged involves a two-stage pipeline: first, pose estimation — the inference of the spatial positions of body parts, e.g., keypoints, from video — followed by the mapping and learning of behavioral classes based on the keypoint-derived kinematic trajectories. While widely adopted, this two-stage pipeline comes with some notable costs. Although pose estimation tools (Mathis et al., 2018; Graving et al., 2019; Pereira et al., 2022) have made high-quality pose estimation accessible, training these models over thousands of video frames represents a significant time sink and requires trained annotators (Sturman et al., 2020). The range of behavioral classifiers built on top of them (Hsu and Yttri, 2021; Segalin et al., 2021; Luxem et al., 2022; Goodwin et al., 2024; Weinreb et al., 2024) also presents a temporal tradeoff; while training behavioral classes is comparatively quicker than training keypoints (Sturman et al., 2020), these classifiers are designed to run offline, downstream of a separate pose-estimation stage, and typically summarize kinematics over temporal windows that can outlast the briefest behaviors of interest. Additionally, these pose-dependent methods inherit a fundamental ceiling: their performance is bounded by the quality of the underlying keypoint tracking.

A qualitatively different approach has recently emerged that bypasses pose estimation entirely, instead performing what has been termed direct video-to-behavior mapping — the inference of behavioral labels from raw video without passing through an intermediate keypoint representation (Skovorodnikov et al., 2025; Gonzales et al., 2026). By learning from the images instead of the keypoints, these methods retain information that keypoints discard, including contextual cues (e.g. food, proximity to in-cage elements), and are resilient to the common failure conditions of pose estimation, e.g., occlusion and low-contrast images. This paradigm utilizes foundation models, large neural networks trained on massive video datasets that learn generalizable spatiotemporal representations. VideoPrism (Zhao et al., 2024), a general-purpose video encoder trained on web-scale video rather than on any behavioral task, outperformed task-specific models on animal behavior benchmarks with a single frozen model. FERAL (Skovorodnikov et al., 2025) built upon this framework with a supervised end-to-end system that fine-tunes V-JEPA 2 (Assran et al., 2025) on user-provided annotations. Similarly, CBAS (Perry et al., 2025) utilized an image foundation model, DINOv2 (Oquab et al., 2024), to achieve efficient, lightweight data processing, enabling continuous segmentation over multiple weeks.

Despite their advantages, these approaches share important limitations. Current video foundation models carry substantial computational costs — requiring significant GPU memory and processing time — so classification must be made retrospectively, offline. While CBAS notably keeps pace with its 10 FPS acquisition rate, it does so in 30-minute batches after the video has been collected. Depth camera-based MoSeq is reported to be configurable to provide 60-100ms latencies approaching real-time (30 FPS acquisition rate), with triggering delays in the hundreds of milliseconds after behavior onset (Markowitz et al., 2023). Although some pose-estimation methods can be configured to process pose in real-time at rates up to 100 FPS (Kane et al., 2020; Schweihoff et al., 2021; Pereira et al., 2022), these methods can only deliver behavior in as much as the behavior can be defined via thresholding a specific pose value, e.g. ‘approach’ as inter-animal distance or ‘reach initiation’ as paw speed. With limited online classification approaches available, researchers are largely prevented from using real-time behavior as a signal for closed-loop experimental control, adaptive experimental paradigms, or continuous monitoring of freely behaving animals over extended durations in which backlogs of unanalyzed video would become prohibitive. It also limits applications in clinical or translational settings where continuous surveillance with triggered perturbations could enable critical inroads (Cavallo et al., 2025; Dixon et al., 2026).

Here we present B-REAL, a supervised machine learning platform that maps raw video directly to behavior with end-to-end inference latencies below 15 ms using only standard GPU or CPU processing power. Because it requires no keypoints, B-REAL inherits neither the annotation burden nor the failure modes of pose estimation. We first benchmarked B-REAL against established pose-based and pose-free methods on mouse social behavior. Using substantially less training data, it performed equivalently to either methods trained on the full benchmark dataset. We then asked whether B-REAL could identify brief, low-occurrence behaviors across a range of fly aggression behaviors lasting as little as a single frame. The platform outperformed both methods in every paired comparison from one minute of labels upward. As an empirical test of its experimental application, we used B-REAL to deliver liquid reward to mice whenever grooming was detected; in this closed-loop experiment, the rate of grooming increased 129% over the course of training. Finally, to leverage the representations learned by video foundation models while preserving real-time inference, we distilled V-JEPA 2 into B-REAL’s initial weights; with this initialization, 30 seconds of labeled video achieved accuracy comparable to 5–10 minutes of labels from an ImageNet initialization. Together, these results demonstrate that behavior, classified in real-time directly from raw video, can serve as a signal for closed-loop experimental control, requiring only minutes of annotation and standard hardware.

## RESULTS

We sought to create a platform in which a behavior could be identified accurately without keypoints and in real-time, defined here as an end-to-end latency shorter than the interval between successive frames, such that no frame goes unclassified as video is acquired. To this end, we adapted the EfficientNet architecture (Tan and Le, 2019) by inserting Temporal Shift Modules (TSM; Lin et al., 2019) between its convolutional stages (Figure 1A). EfficientNet is a family of accurate, computationally lightweight convolutional neural networks, originally designed for fast inference on mobile phones (Tan et al., 2019; Tan and Le, 2019). It has proven broadly useful across computer vision tasks, including as a backbone for animal pose estimation (Mathis et al., 2021), while maintaining fast inference on both powerful GPUs and compact edge devices. TSM, developed for video action recognition, allows an image network such as EfficientNet to capture motion by exchanging information between neighboring frames, without adding parameters or computation (Lin et al., 2019). Importantly, in order to achieve rapid classification, this approach avoids the training or use of keypoints entirely, and therefore inherits neither the annotation burden of pose estimation (Sturman et al., 2020) nor its characteristic failure modes.

**Fig. 1.**
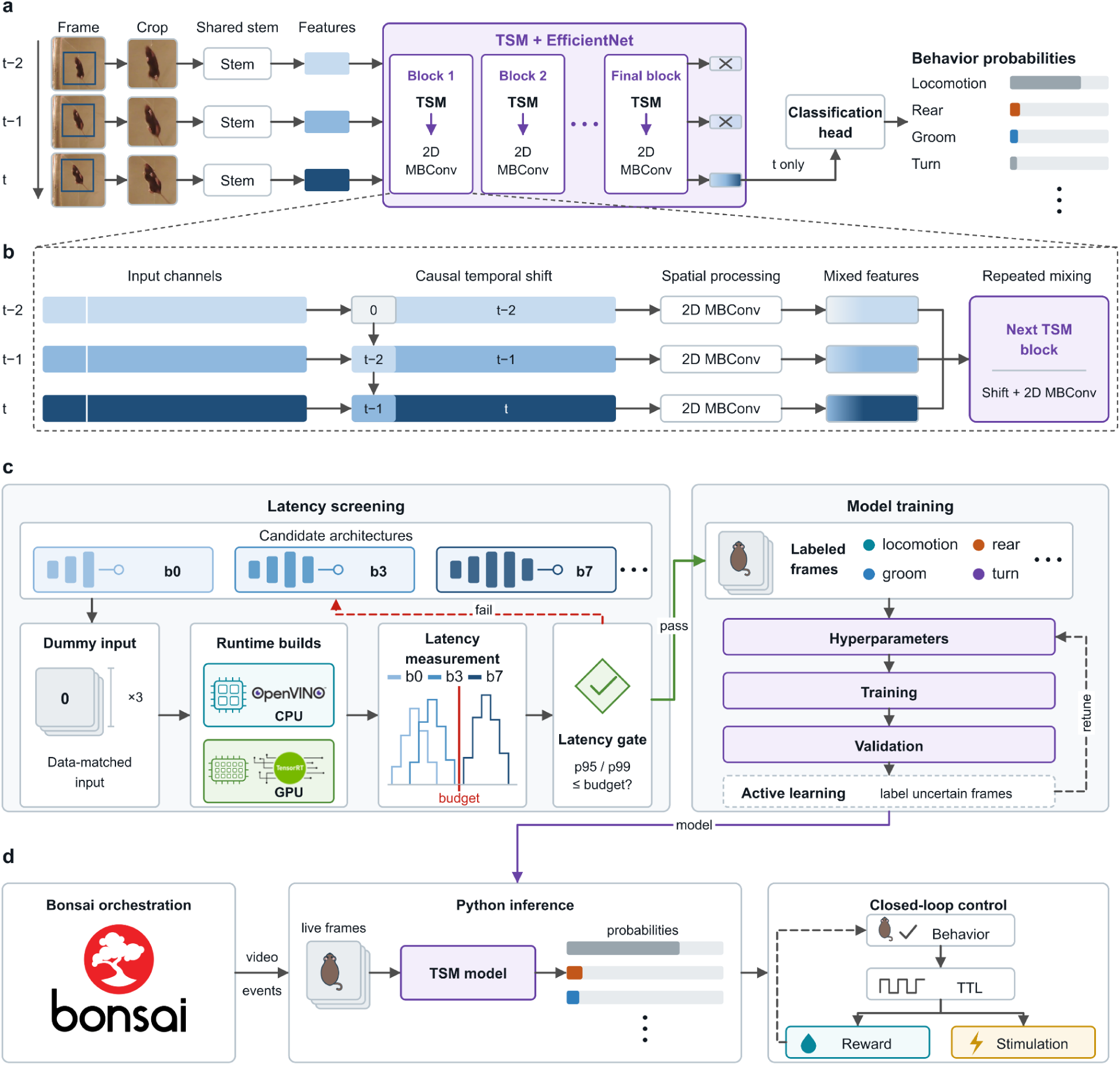
B-REAL: a pose-free behavior classifier with latency screening for closed-loop deployment. **a**, Real-time classification. At each frame the animal is cropped from the video. The crops for frames t−2, t−1 and t pass through a shared stem into a stack of TSM + EfficientNet blocks. Only features from the last block of the current-frame stream reach the classification head, which outputs one set of behavior probabilities per frame. **b**, An enlarged view of one TSM + EfficientNet block from **a**. A fraction of channels from each frame is replaced by the same channels from the previous frame, padding the first frame with zeros. This mixes temporal context with no additional learned parameters. Repeated mixing across blocks enables learning of complex spatiotemporal features. **c,** Model development workflow. Candidate EfficientNet backbones (b0/b3/b7 shown) are screened for latency before any training. A dummy input matched to the data’s shape is run through OpenVINO (CPU) and TensorRT (GPU) builds, and only architectures whose 95th- or 99th-percentile latency fits within the per-frame budget are trained. Training then optionally iterates, using active learning on uncertain frames. **d**, Closed-loop deployment. Bonsai streams video and events to the Python inference process. The behavior probabilities may then drive a TTL which delivers reward, stimulation, or changes in context.

Briefly, a sequence of frames—a ‘clip’—capturing individual or multi-animal behavior is collected and optionally cropped around the subject of interest. A basic set of features, such as detected lines and edges, is obtained separately per-frame using the frozen convolutional stem from the base layer of EfficientNet, and these features are then passed into a sequence of TSM-convolutional blocks. Within each block, a small subset of the within-frame features is shifted across frames before being processed by the next convolutional layer (Figure 1B), such that across many blocks the model learns to extract features that are both spatially and temporally meaningful as spatial information within each frame is mixed and reprocessed across frames. These convolutional layers are initialized with EfficientNet weights pre-trained on the ImageNet dataset (Deng et al., 2009) but are fine-tuned during learning. Notably, the shift operation carries no learned parameters and adds no multiply-accumulate operations, so temporal context is acquired at essentially no cost in either model size or inference time—the property that makes the architecture tractable for real-time use.

Because we aim to classify the subject’s behavior specifically at the time of the current frame (frame t, Figure 1A), we deviate from the original TSM approach by selecting the features of only the last frame in the clip for classification, rather than averaging them across frames. Through the temporal shifts, these features already incorporate information from the preceding frames, which provide context to capture the temporal domain of the behavior; this is useful for classifying behavioral bouts that may last for a shorter duration than the clip context, or in the practical case in which a clip contains a behavioral transition. Because no later frame is involved, the prediction for frame *t* depends only on that frame and those before it as real-time use requires (channels that would receive information from a later frame are filled with zeros). We then process the output features with a lightweight classification head which produces a set of output pseudo-probabilities across the selected classes.

Deploying such a model in a closed loop requires knowing, before committing to training, whether it will run fast enough; we therefore provide a latency screening step (Figure 1C). This screening approach is possible because processing time is a function of the structure of the network, not of the precise values of its weights. Users first select a candidate architecture from amongst the eight EfficientNet backbones, b0-b7, where the optimal choice is generally the largest possible model which meets the compute budget, e.g. the desired frame rate. B-REAL generates a set of dummy-inputs that match the specifications of the experimental setup, including the number of frames per clip and their sizes, and packages the candidate architecture into an efficient format for inference—either TensorRT for Nvidia GPUs or, notably in the case that a GPU is not available, OpenVINO for Intel CPUs. A distribution of latencies is then computed and its 95th or 99th percentile compared against the described budget, which is 16.7 ms for 60 FPS acquisition and 33.3 ms for 30 FPS. We use a high percentile rather than the mean because in a closed loop every frame must be processed on time: an architecture whose average latency fits the budget can still overrun it on occasional slow frames, delaying or skipping those classifications, whereas clearing the 95th or 99th percentile ensures that no more than 5% or 1% of frames exceed it. The largest model architecture which still clears the budget is then trained and optionally hyperparameter-tuned; if test metrics indicate the trained model is lacking, one can iteratively label additional frames and retrain until sufficiently accurate. To aid in producing behavioral labels, we provide a simple graphical user interface that loads videos directly and supports efficient frame-precise labeling across behaviors, removing the need for separate labeling software such as BORIS (Friard and Gamba, 2016).

The trained model can be exported to TorchScript for direct execution within custom Python control scripts, or into a Bonsai-accessible format (Lopes et al., 2015) (Figure 1D). Bonsai is a widely used visual coding framework that aids in the deployment of multi-tiered experimental paradigms, supplying frames and hardware events to the inference process while the resulting behavior posterior drives a TTL line capable of gating a reward, a sensory stimulus, or an optogenetic manipulation. We provide an example workflow which enables real-time data acquisition and inference, and were able to readily achieve 30 FPS processing speed with an NVIDIA 1080 Ti GPU. Importantly, even without the real-time application, B-REAL achieves the major advance of providing behavior classification without the effort-intensive training of pose estimation keypoints.

### Rapid, accurate social behavior classification with minimal training

We first evaluated our framework on the CalMS21 benchmarking dataset (Sun et al., 2021) of mouse social behavior, labeled per-frame as representing attack, investigation, mounting, and a catch-all ‘other’ class. We compared our method against two established supervised classification approaches, A-SOiD (Tillmann et al., 2024) and CBAS (Perry et al., 2025). A-SOiD is an offline, keypoint-based classifier that uses active learning to reach high accuracy from few labels and achieves rapid classification (>100 fps equivalent). CBAS is also pose-free, learning an LSTM on top of features obtained from a frozen image foundation model; it classifies at acquisition rate but only in 30-minute batches after the video has been collected. We note that neither of these approaches were designed for the combination of real-time use at a high temporal resolution; instead, each represents the state of the art in offline and in batched classification. Unless otherwise noted, B-REAL and CBAS were trained and evaluated on identical frames, cropped around both mice using the keypoint estimates provided with CalMS21 (see Methods). On held-out test video, B-REAL’s per-frame predictions closely follow the ground-truth annotation, whereas A-SOiD’s changes more slowly, reflecting the time bin over which it summarizes keypoint trajectories (Figure 2A) and a fundamental limitation of most keypoint methodologies.

**Fig. 2:**
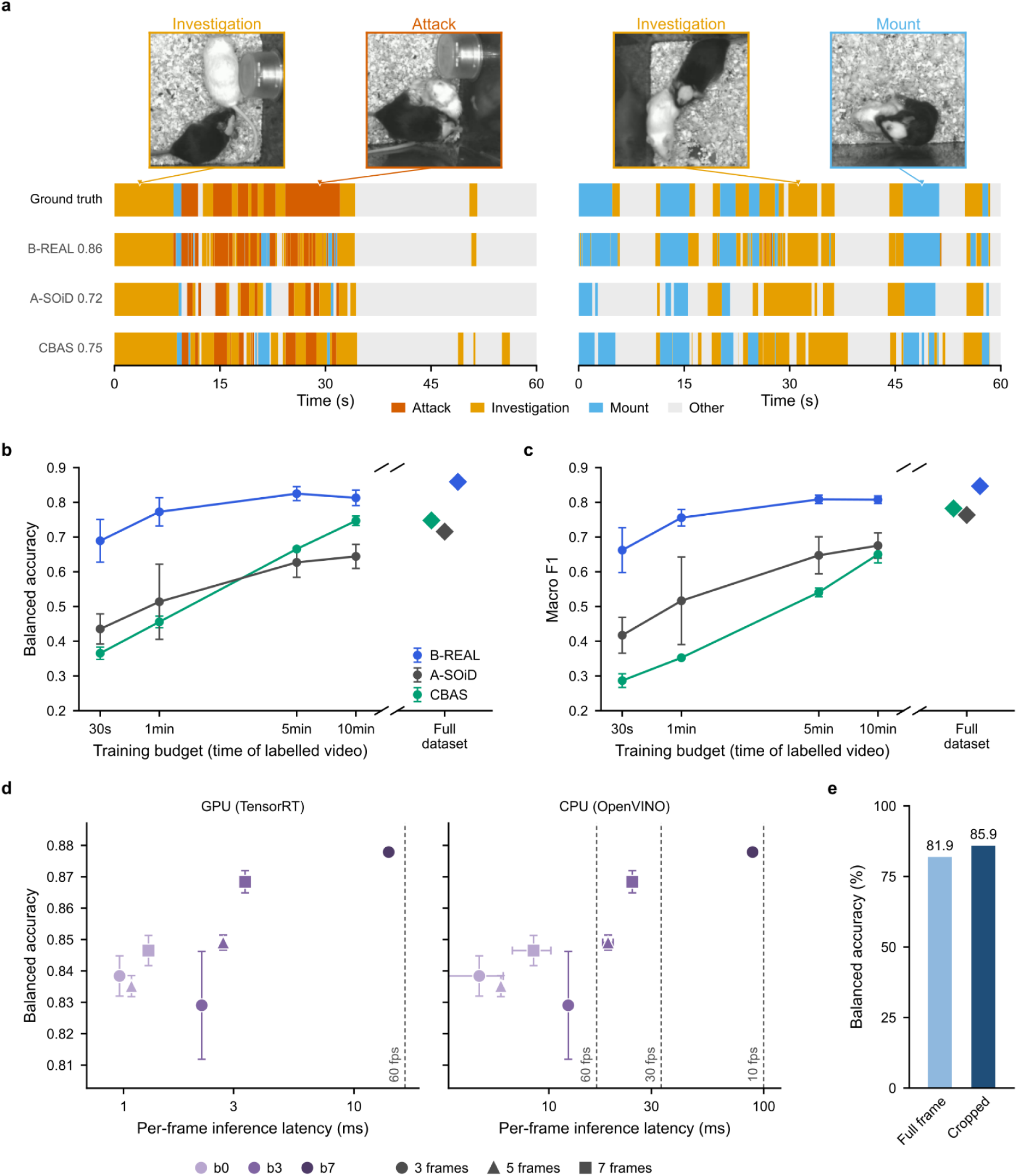
B-REAL enables real-time classification of social behavior with state-of-the-art performance. **a**, Two example held-out CalMS21 test windows. Top, example frames at the marked times reflecting resolution of the video data. Below, the per-frame class over 60 s from ground truth, B-REAL, A-SOiD and CBAS. The balanced accuracy beside each method is computed over the full 19-video test set. **b**, Test-set balanced accuracy as a function of the amount of labelled training video, mean ± s.d. (B-REAL, 8 independent label draws; A-SOiD and CBAS, 3 seeds). Right, single runs on the full training set. **c**, Same as **b** but for macro F1 over the four classes, from the same runs. **d**, Balanced accuracy against per-clip inference latency for EfficientNet b0/b3/b7 backbones (colour, light to dark) on 3-, 5- and 7-frame clips (marker shape). Latency was timed on an RTX 5090 GPU (TensorRT, left) and an Intel CPU (OpenVINO, right). Accuracy is mean ± s.d. over seeds (b7, single seed), and latency is mean ± s.d. over 7 launches. Dashed vertical lines indicate the per-frame budgets at 60 fps (16.7 ms), 30 fps (33 ms) and 10 fps (100 ms). b7 with 5- and 7-frame clips are omitted. **e**, Balanced accuracy with full-frame and animal-cropped input.

To assess the sample efficiency of our method, we compute the balanced accuracy (the average of per-class recall scores) on the test set as a function of the quantity of training data used to train each model (Figure 2B). At every timed budget, B-REAL outperforms both methods substantially (Welch’s t-test, Holm-corrected p < 0.05 in all eight comparisons). The performance disparity is largest at the smallest budgets: with just 30 seconds of labeled data, B-REAL exceeds A-SOiD by 25 balanced-accuracy points and CBAS by 32. Importantly, with one minute of randomly-sampled labeled data, B-REAL surpasses both A-SOiD and CBAS trained on the full CalMS21 dataset, by 5.7 and 2.5 points respectively. Because balanced accuracy does not penalize false positives, we also scored each method by macro F1, the average of the per-class F1 scores, and found the same ordering (Figure 2C). B-REAL leads at every budget, by 24.5 and 37.6 points over A-SOiD and CBAS at 30 seconds (Welch’s t-test, Holm-corrected p < 0.05 in seven of eight comparisons; the exception is A-SOiD at 1 minute, p = 0.079, limited by that method’s three-seed variance). On this stricter measure, B-REAL trained on five minutes of data surpasses A-SOiD and CBAS trained on the whole dataset by 4.5 and 2.6 points, respectively. B-REAL achieves state-of-the-art performance in social behavior classification with only limited amounts of training data.

To assess the extent to which performance improves with larger model backbones and longer clip contexts, we trained multiple versions of B-REAL on a fixed 20-minute labeling budget with various hyperparameter combinations (Figure 2D). For three EfficientNet backbones—b0, b3, and the largest available, b7—we trained models with 3-, 5-, and 7-frame clips as context. Within a given backbone, longer clips generally improved performance at the cost of inference time; this is most evident with the b3 backbone, which shows performance rising from 82.9% to 86.8% balanced accuracy as the clip grows from 3 to 7 frames (Figure 2D). The best reproducible configuration, b3 with a 7-frame clip, reaches 86.8% balanced accuracy for 3.5x the per-clip GPU latency of the smallest model; a single run of the much larger b7 backbone at a 3-frame clip edged this to 87.8% but at 14.7x the latency. Notably, every b0 and b3 configuration runs faster than the 30 FPS acquisition rate on both our workstation GPU and an Intel CPU; the slowest of them, b3 with a 7-frame clip, sustains 297 FPS on GPU and 41 FPS on CPU. The b7 backbone remains above acquisition rate on GPU at a 3-frame clip (71 FPS).

Finally, to assess the contribution of cropping, we retrained B-REAL on full, uncropped frames with all other settings unchanged, and found that cropping improved performance (Figure 2E). Because every frame must be resized to the fixed input size of the network, much of the high-resolution detail specific to the animal is lost to accommodate the full image. Cropping out the empty arena fills the input mostly with the animal, preserving fine details such as paw and head movements that would otherwise be lost to down-sampling. This improvement comes at the expense of context, as a crop centered on the animal discards its surroundings and its position in the arena, and with them the translational movement that may help identify behaviors defined by location or movement. While there is a several point increase in accuracy with cropping, for users wanting to incorporate the full scene, B-REAL performance with the full frame still outperforms the other approaches shown here. Where keypoint estimates are not available, as in our closed-loop experiments and pre-training corpus (see Figure 5), B-REAL instead locates the animal by blob detection (see *Methods*). Our instantiation involved a simple background subtraction step followed by morphological operations to locate the center of mass of the mouse in every frame.

### B-REAL detects rare and brief behaviors from only minutes of labeled data

While the social actions evaluated in CalMS21 persist across extended frame sequences, we next asked whether B-REAL’s direct spatiotemporal processing could identify rapid, low-occurrence behaviors in a second species. To do so, we tested it on the aggression subset of the Fly-vs-Fly dataset (Eyjolfsdottir et al., 2014). This benchmark features ten 30-minute videos captured at 30 FPS of pairs of male *Drosophila*, annotated per fly and per frame for five aggressive behaviors (lunge, wing threat, charge, hold, and tussle) alongside a catch-all ‘other’ class. Class prevalence varies across more than two orders of magnitude, with median bout lengths spanning from a single 33 ms frame to over 400 ms (Figure 3A). The charge category represents an extreme on both axes, comprising just 0.016% of scored fly-frames; 56% of its 117 labeled bouts persist for only a single video frame (33 ms), and none exceed four frames. Lunge, the second-briefest action (median duration 133 ms), occupies 70 times as many frames. Figure 3B illustrates sequential frames capturing individual bouts of charge, lunge, and wing threat.

**Fig. 3.**
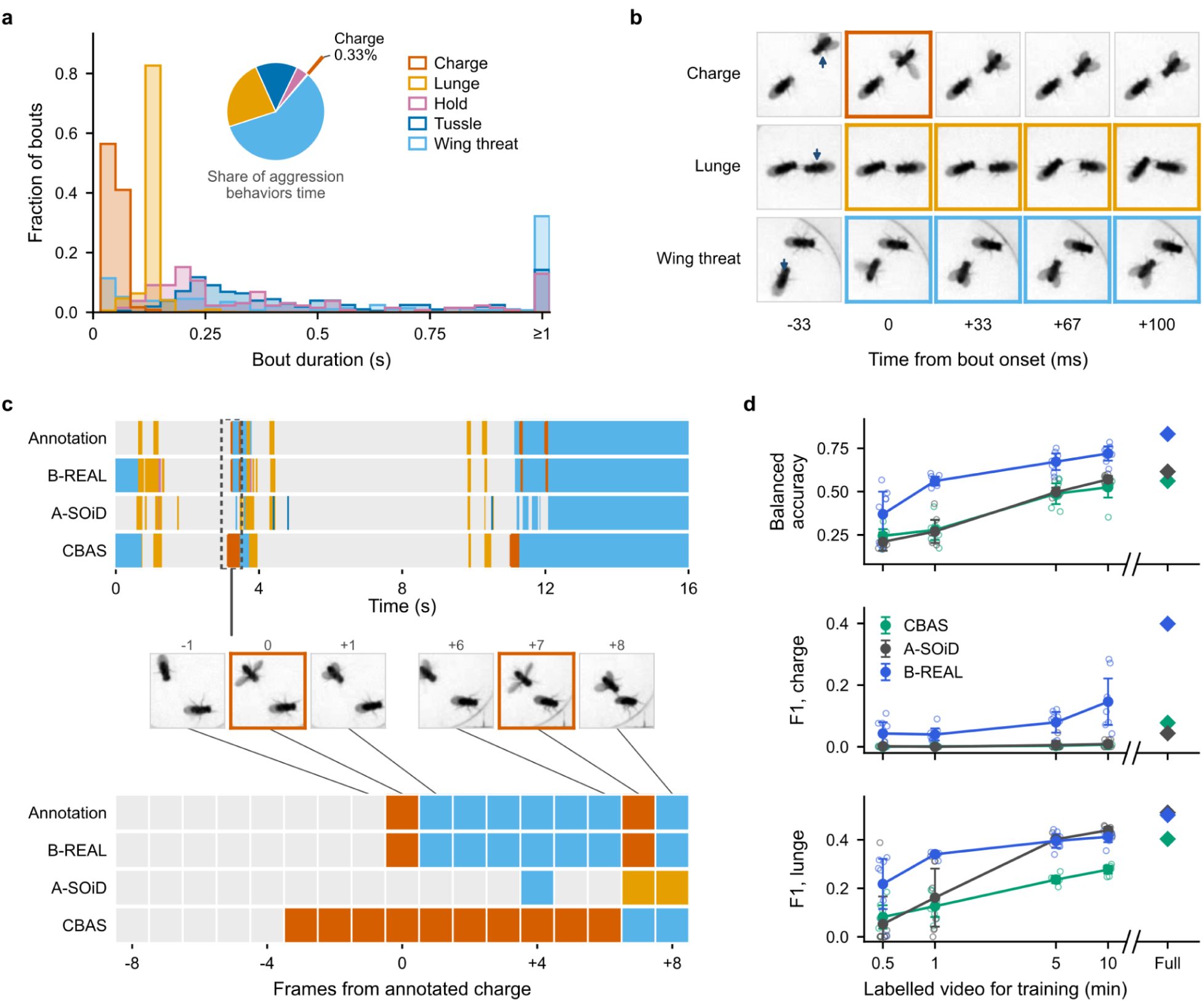
B-REAL identifies a rare, brief fly social behavior with uniquely strong accuracy. **a**, Annotated bout durations for each behavior over all ten movies, as a fraction of that behavior’s bouts. There is one bin per video frame (33 ms), and bouts of 1 s or longer are pooled in the last bin. Inset, share of aggression time spent in each behavior; aggression time is the 4.8% of scored fly-frames in which any of these behaviors occurs. Charge (pulled out) is 0.33% of aggression time and 0.016% of all frames. 56% of charge bouts last a single frame, and none exceeds four. **b**, Five consecutive frames (30 fps) around one annotated bout each of charge, lunge and wing threat, aligned to onset (0 ms). Outlined frames are annotated, and the arrowhead marks the fly performing the behavior. All five behaviors are shown in Supplementary Figure. S1. **c**, Top, per-frame class over 16 s of held-out test movie 8, the test set’s densest cluster of charges (chosen from the annotation alone). Rows show the annotation, then B-REAL, A-SOiD and CBAS trained on the full training set; charge frames are drawn at a minimum width. In this window B-REAL detects all 4 annotated charge bouts (a charge call within ±1 frame) with no false alarms; A-SOiD detects none, and CBAS detects 2 with one false alarm. Bottom, the 17 frames around the first charge (dashed box), one cell per frame, with the video frames before, at and after each annotated charge. **d**, Balanced accuracy, charge F1 and lunge F1 on the held-out test movies against labelled training video (mean ± 95% CI over 8 label draws per budget, the same draws for all three methods; individual draws in the background). Right, single runs on the full training set. From 1 min upward, B-REAL beats both baselines in all 8 paired draws on balanced accuracy and on charge F1 (Wilcoxon *p* = 0.008); at 30 s neither lead is reliable. Lunge F1 favours B-REAL at 30 s and 1 min only. F1 is shown rather than recall because at 10 min CBAS matches B-REAL’s charge recall with a precision of 0.003.

We implemented an identical three-way benchmark to our CalMS21 pipeline. Each model variant was trained on matching label draws across five training videos from the official split and scored against per-frame ground truth across five held-out evaluation videos. B-REAL utilized our most basic configuration (Figure 2): the simplest EfficientNet-b0 backbone with a 3-frame clip window and ImageNet initialization, operating on uncropped frames without altering default hyperparameters. A-SOiD was trained using five tracking keypoints per fly (head, centroid, tail, and both wings) derived from native ellipse-and-wing trajectories; following published guidelines, its temporal bin was aligned to the briefest action of interest (one frame), as the default 100 ms bin discards 39% of charge annotations prior to optimization. CBAS was evaluated on the raw video stream. Training allocations ranging from 30 seconds to 10 minutes were sampled across eight random draws, supplemented by a single evaluation on the complete training set.

B-REAL consistently demonstrated superior sample efficiency (Figure 3D). From one minute of annotated video upward, it outperformed both baselines on test-set balanced accuracy in all eight paired draws at every budget (Wilcoxon signed-rank test, Holm-corrected p = 0.016, the minimum attainable with eight draws). At one minute, B-REAL reached 0.56 balanced accuracy, compared to 0.27 for A-SOiD and 0.28 for CBAS. At 30 seconds, its mean advantage (0.37 vs. 0.21 and 0.24) was not consistent across draws (6/8 and 5/8 wins; p = 0.16 for both). Given five minutes of labeled data, B-REAL (0.67) surpassed both baselines trained on the entire training set (0.62 for A-SOiD, 0.56 for CBAS), reaching 0.83 when itself trained on the full training set.

This performance differential was most pronounced for the rarest, briefest behavior in the dataset—charging. To account for false positive rates on rare classes, we evaluated per-class F1 scores. B-REAL’s charge F1 score exceeded both baselines in all eight draws at 1, 5, and 10 minutes (p = 0.016); on the full dataset, its F1 score (0.40) was five-to nine-fold higher than A-SOiD (0.04) and CBAS (0.08). The comparison architectures exhibited distinct failure modes: A-SOiD generated virtually no positive charge predictions (0.004 recall at 10 min), whereas CBAS matched B-REAL’s recall (0.48 vs. 0.47 at 10 min) but at a high false-positive cost (0.003 precision vs. 0.10 for B-REAL). Frame-level predictions illustrate this contrast directly (Figure 4C): across the test set’s densest charge cluster, B-REAL predicted a charge within one frame of all four annotated events without false positives, whereas A-SOiD detected zero bouts and CBAS identified two alongside a false positive.

**Fig. 4.**
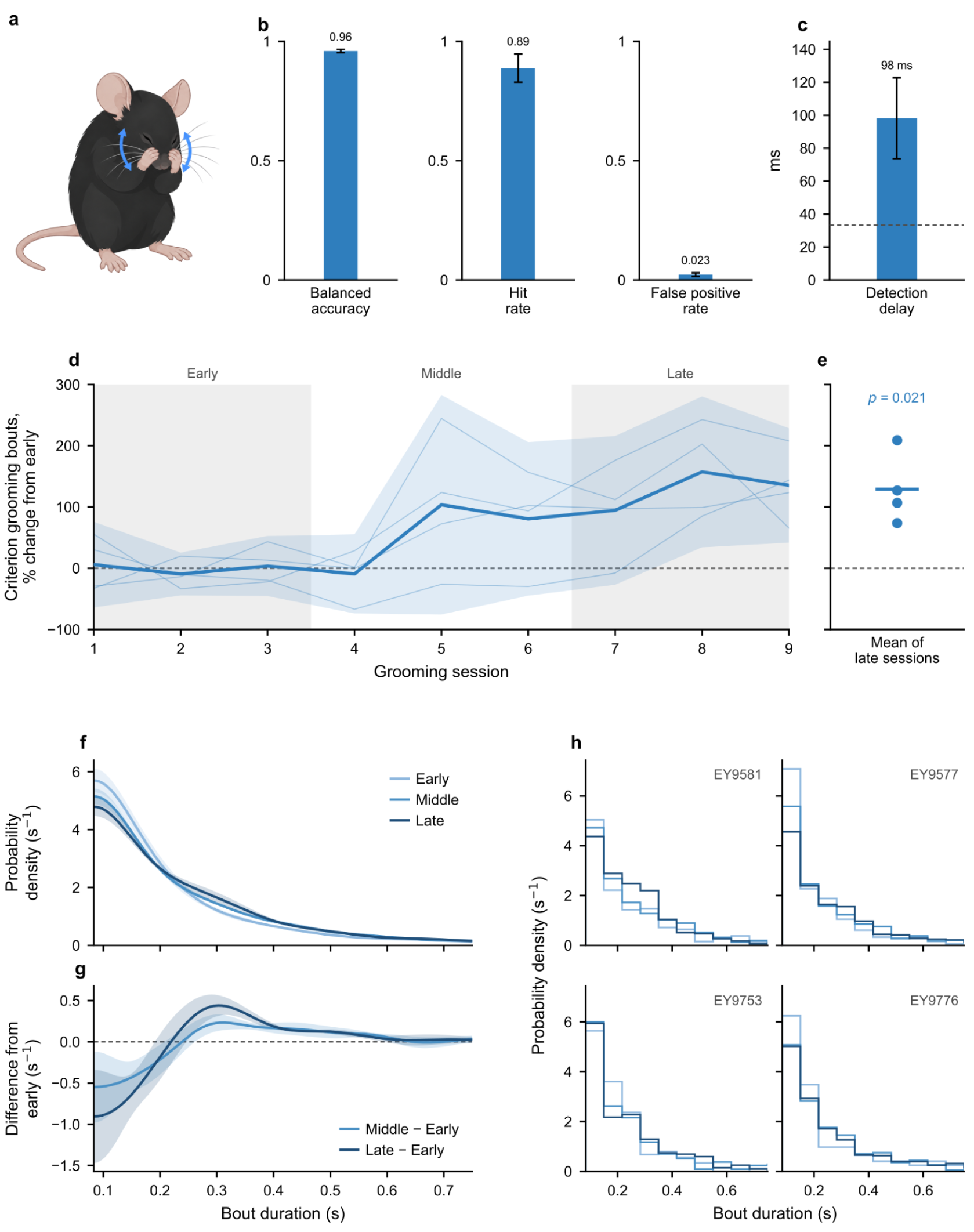
A real-time grooming classifier drives increases in grooming bout rates. **a**, Head grooming, the rewarded behavior, involves a repetitive movement of the forepaws across the face and head of the animal. **b**, Held-out performance of the real-time grooming classifier: balanced accuracy, hit rate and per-frame false positive rate, each on its own 0–1 axis; mean ± s.d. over three training seeds. **c**, Delay from grooming onset to first detection, mean ± s.d. over the same seeds (98 ms). The dashed rule is one frame at 30 FPS (33 ms). **d**–**h,** Mice (n = 4) received sweetened water when the online classifier detected grooming on three consecutive frames (100 ms). A criterion grooming bout is a run of at least three consecutive frames classified as grooming, which triggered reward during training. Early, middle and late are grooming-block sessions 1–3, 4–6 and 7–9. **d**, Rate of criterion grooming bouts in each session, as % change from each animal’s own early sessions. These sessions were already rewarded, so they are not a pre-reward baseline. Thin lines, individual animals; thick line, mean; band, 95% CI across animals; grey shading, the early and late windows. **e**, Mean change in the late sessions (dots, animals; line, mean of animals): +129% (paired t-test on per-animal means against the early sessions, p = 0.021. **f**–**h**, Duration of criterion grooming bouts in the early, middle and late sessions (4,408 bouts in 36 sessions). **f**, Probability density (mean and s.e.m.) of grooming bout duration across animals (animals weighted equally). Gaussian kernel (σ = 50 ms). The x axis starts at the shortest possible bout (100 ms). **g**, Middle and Late densities minus Early, differenced within each animal. Dashed line = no change. **h**, Per animal effects in two frame (67ms) bins, in the same density units as f. Bouts shift slightly from the shortest durations toward 0.25–0.5 s.

B-REAL’s performance advantage concentrates primarily on high-speed actions and minimal annotation budgets. For lunge, B-REAL’s F1 score surpassed A-SOiD at 30 seconds and 1 minute (+0.17 and +0.18), though A-SOiD converged to comparable accuracy from 5 minutes onward (−0.01 and −0.03 at 5 and 10 min; p = 0.55 and 0.078); B-REAL maintained higher scores than CBAS across all eight draws from 1 minute upward. On longer-duration actions (hold, tussle, and wing threat, median durations 8–13 frames), A-SOiD achieved higher F1 scores at 10 minutes (0.47, 0.58, and 0.78, versus 0.43, 0.49, and 0.61 for B-REAL). Consequently, A-SOiD’s macro F1 score edged B-REAL at 5 and 10 minutes (0.50 vs. 0.48, and 0.54 vs. 0.51), though its balanced accuracy remained lower. These results indicate that pose-based pipelines remain effective for sustained behaviors given sufficient training data, whereas B-REAL offers a critical advantage for closed-loop applications: precise detection of brief, transient behavioral events under low-label regimes.

### Real-time closed-loop manipulations with B-REAL can shape naturalistic behaviors

To assess the capacity of our model to drive behavioral changes in a closed-loop setting, we trained mice to associate the performance of grooming with a water reward (Figure 4). Grooming is an important behavior in natural ethology and the study of psychiatric disease, including OCD, anxiety, and autism (Kalueff et al., 2015). We used the smallest EfficientNet backbone and a three-frame clip size. The resulting model achieved a balanced accuracy of 0.96 and recalled 89% of grooming frames on a held-out test set, at a per-frame false positive rate of 2.3% (Figure 4B), suggesting that the platform would be sufficient to detect and shape behavior. We also computed the average delay between the first frame of a bout and the first predicted instance, which was ∼3 frames (98 ms; Figure 4C). This delay for behavior onset between the human rater and B-REAL in part reflects the common observation that a substantial proportion of the disagreement between human raters occurs at the transition into and out of a behavior (Segalin et al., 2021). Given this known ambiguity in establishing when a behavior truly starts, B-REAL provides an option to down-weight the noisy bookends of a behavior bout. Here, and in all instances noted previously in the paper, we utilize this focused-training option - while still retaining high accuracy with the human-rater ‘ground truth’. Additionally, we assumed that to be considered sufficient for triggering, a bout would require a criterion number of consecutive, positively identified frames; and as such, a small delay in initial detection would be moot - particularly in light of the high hit rate (Fig 4B), i.e. the ability to positively identify some part of each bout. We test these functional assumptions for reinforcing naturalistic behaviors with closed-loop rewards below.

We trained mice (n=4) to receive a water reward for performing head grooms. Upon the detection of three consecutive head grooming frames (100 ms), an audible solenoid opened to deliver sweetened water through a spout in the wall on one side of an arena; a light above the spout also turned on and stayed on until the spout was visited. During the Early phase (sessions 1-3), mice were naive to both the reward mechanisms - e.g. solenoid, light, presence of water - and the specific groom-reward contingency. We introduced both elements together, after multiple arena acclimation sessions, to capture the full progression of learning from naive to trained (see Supplementary Figure S2 for reward receipt latencies).

In this rewarded context, mice progressively increased the performance of grooming behaviors compared to their first rewarded sessions, over nine sessions in which grooms elicited reward (Figure 4D). After scoring every session offline, we found that the mean rate of criterion-clearing (≥ 100ms) detection events increased 129% between Early and Late sessions (paired t-test across the four per-animal means, p = 0.021; Figure 4E). In addition to changing the rate of occurrence, reinforcement can alter how behaviors are performed (Hodge & Yttri, 2025). Across the 4,408 criterion bouts in sessions 1–9, the distribution of grooming bout durations progressively shifted from shorter bouts toward bouts of roughly 250-500 ms, presumably reflecting a strategic increase to insure above-criterion grooming durations. The share of bouts lasting 333–500 ms rose in all four animals (+4.1% paired t-test, p = 0.006; p = 0.025 after Holm correction over the four duration bands examined, see Methods), and the distribution above 0.5 s was unchanged. The onset of the reward light and sound cues upon detection likely limited the extent of these duration changes. Together these results demonstrate that closed-loop reinforcement delivered by B-REAL is fast enough and specific enough to increase the rate of a naturalistic behavior, using a detection criterion of only three frames.

### Additional V-JEPA 2 pretraining on unrelated video can improve label efficiency with no inference cost

We next explored whether B-REAL’s label efficiency could be improved further by initializing it from the representations of a video foundation model. By default, B-REAL’s EfficientNet backbone is initialized from ImageNet weights, which are optimized for static images. Although the TSM mechanism exchanges features across the frames of a clip, those features are never trained to encode motion. Temporal structure must therefore be learned from the labelled examples alone, which is most limiting when labels are scarce. Video foundation models such as V-JEPA 2 (Assran et al., 2025), in contrast, learn generalizable spatiotemporal representations from large video collections and have been used for direct video-to-behavior mapping elsewhere (Skovorodnikov et al., 2025). Running such a model directly, however, precludes real-time operation, as its large vision transformer over 16-frame clips costs far more computation per prediction than B-REAL’s backbone. We reasoned that distilling V-JEPA 2’s representations into B-REAL’s backbone using unlabelled mouse video might provide superior pre-training weights for clip-level behavioral identification while leaving the deployed architecture, and therefore its inference speed, unchanged. Distillation trains B-REAL’s own backbone to reproduce the foundation model’s representations. Specifically, we trained B-REAL to match the spatial-temporal features produced from V-JEPA 2. The V-JEPA 2 network processed a 16-frame sequence into a 1024-dimensional embedding vector, while the smallest B-REAL variant (b0 backbone, 3-frame clip) was optimized to match this target representation using only the final three frames of the sequence. Crucially, the foundation model parameters remained static and target vectors were pre-computed offline, requiring only a single pass through the dataset. By anchoring both clips to the final frame, the target vector represents the immediate behavioral state rather than an average across the preceding time window. This self-supervised approach requires no human annotations, allowing pre-training across arbitrary, unlabeled video files.

The pre-training corpus combined five diverse mouse behavior datasets, providing a broad range of experimental environments: CalMS21 (overhead social interactions; Sun et al., 2021), CRIM13 (side-view cage recordings; Burgos-Artizzu et al., 2012), MABe22 (overhead group behavior on bedding; Sun et al., 2023), a lockbox foraging paradigm (overhead single animal; Reiske et al., 2025), and JAX open-field trials featuring pigmented and albino strains (Geuther et al., 2019). All videos were standardized to 30 FPS to maintain consistent temporal spans across datasets, and subjects were isolated via automated background subtraction bounding boxes. Because feature extraction relies entirely on the frozen foundation model, no behavioral labels were required during dataset assembly. To ask what this diversity contributes, we also distilled a second student from unlabelled CalMS21 training video alone, and compared the two throughout. Importantly, none of these videos were specific to the target data - and in the cases of different colored coats or sideview video, were remarkably different. Additionally, while most of the teacher videos were grayscale, the target video was in color.

Representation distillation substantially restructured feature space prior to fine-tuning. Linear probes trained on frozen backbones across 40 training videos demonstrated that distillation increased test set balanced accuracy on CalMS21 from 0.62 (ImageNet initialization) to 0.71–0.75 across twelve pre-training epochs. Because the linear readout was evaluated on static backbones, this improvement reflects enhanced linear separability of behavioral classes within the feature space. Further analysis confirmed that ridge regression mapped ImageNet features to V-JEPA 2 representations with only R² = 0.47, indicating that over half of the temporal feature variance captured by the video foundation model is absent from standard ImageNet initializations.

When fine-tuned on labelled data, the student distilled on CalMS21 video outperformed the ImageNet initialization at every budget, with the largest gains under extreme data scarcity. On CalMS21 Task 1, it increased test balanced accuracy from 0.69 to 0.80 with just 30 seconds of training data (+11 percentage points, 95% CI [6.3, 15.9], 8/8 draws), from 0.77 to 0.83 at 1 min (+6 [3.8, 8.5], 8/8), from 0.83 to 0.86 at 5 min (+3% [1.1, 5.1], 7/8) and from 0.81 to 0.85 at 10 min (+ [1.3, 5.5], 8/8). To control for potential frame-level overlap, we repeated the sweep with video-disjoint draws, in which each draw’s labelled frames came only from its own group of training videos (Figure 5B; n = 5 groups). The gain held at low budgets (+11 at 30 s, 95% CI [5.2, 16.6]; +9 [5.1, 13.3] at 1 min; 5/5 groups each) but was not resolvable at 5 min (+4 [−3.6, 12.6]) or 10 min (+3 [−4.1, 9.1]) with five groups. Distilling from the five-source corpus instead gave smaller gains that did not clear zero at any budget (+5 [−2.0, 11.6] at 30 s; +2 [−2.2, 6.4] at 1 min; +2 [−1.4, 5.4] at 5 min) and reversed at 10 min (−3 [−9.0, 2.9], 1/5 groups). On the dataset it was evaluated on, then, the broader corpus was no better, and at larger budgets possibly worse, than pre-training on that dataset’s own unlabelled video. This improved sample efficiency dramatically lowers the manual annotation burden. B-REAL trained on a mere 30 seconds of labeled video (0.80) achieved parity with ImageNet-initialized models trained on 5 (0.825, p > 0.5) or 10 minutes (0.813, p > 0.5) of labeled data, representing a ten-to twenty-fold reduction in required human annotation time.

**Fig. 5.**
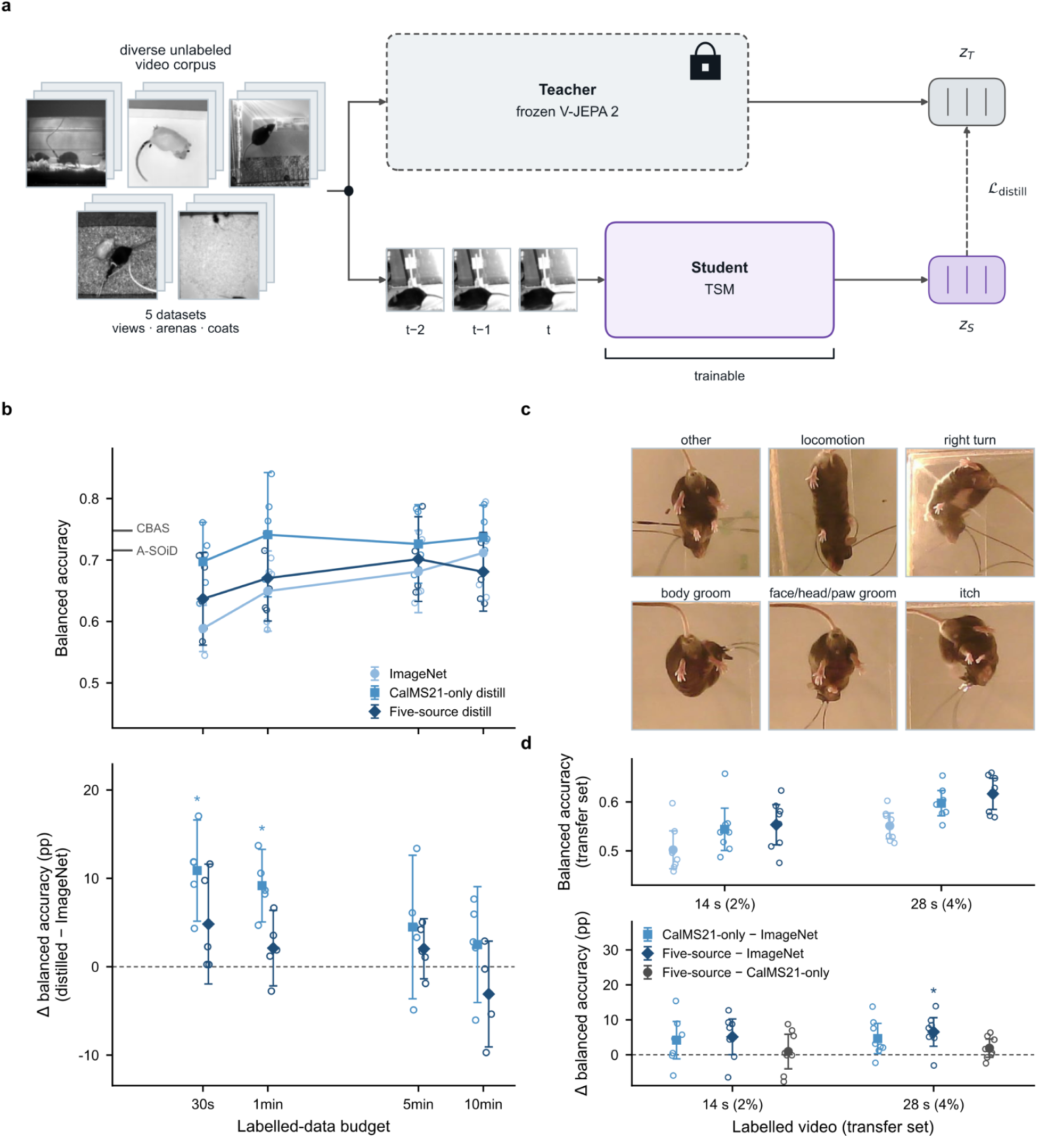
Distilling a frozen video foundation model into B-REAL improves label efficiency. **a**, Pre-training setup. A frozen V-JEPA 2 teacher and the TSM student used at deployment (plus a projector) embed the same unlabelled clip, and a distillation loss encourages the student’s features to match those of the teacher. The student’s weights then serve as the initialization for fine-tuning for real behavioral classification tasks. **b**, Top, CalMS21 Task 1 balanced accuracy against labelled-data budget for B-REAL with three starting points: ImageNet weights, a student distilled on unlabelled CalMS21 training video, or a student distilled on the five-source corpus shown in **a** (mean ± 95% CI over 5 video-disjoint label draws per budget; individual draws in the background). Grey marks at left show the best balanced accuracy A-SOiD (0.72) and CBAS (0.74) reach on the same test set with the full training set (single runs; Figure 2). Bottom, paired differences from ImageNet for each draw. The CalMS21-only student gains +11 percentage points at 30 s and +9 at 1 min (CI excludes 0; 5/5 draws). The five-source student’s gain never excludes 0 and reverses at 10 min (−3 points; 1/5 draws). **c**, The transfer set is filmed from below, a view absent from every pre-training corpus. Shown is one frame from each of its six scored classes as the network sees it: the middle frame of that class’s longest annotated bout. **d**, Transfer-set accuracy with 14 s and 28 s of labelled video (8 label draws per budget; key as in **b**). Here the CalMS21-only student is subsampled to the same number of anchor clips as the five-source corpus, so the two differ only in diversity. Both beat ImageNet at 28 s, and the difference between the five-source and CalMS21-only students includes 0 at both budgets. Stars **b**–**d**): one-sample t-test of the per-draw paired differences against 0, uncorrected (* p < 0.05, ** p < 0.01), so a star marks exactly the CIs that exclude 0.

We further tested whether these feature representations generalize to novel recording setups by fine-tuning on an unseen single-animal dataset captured from a bottom-up ventral perspective. Crucially, no ventral view recordings were included in any pre-training dataset: four of the five corpus sources are overhead and the fifth is side-on, so no pre-training clip contains a ventral view of a mouse. At minimal training allocations (14 s and 28 s of labeled video, both below the smallest CalMS21 budget), both pre-trained initializations outpaced ImageNet. Given 28 seconds, balanced accuracy rose from 0.55 to 0.60 with a CalMS21-only corpus (+5 percentage points, 95% CI [0.3, 9.0]) and to 0.62 with the five-source corpus (+7 [2.4, 10.6]), with the same ordering at 14 seconds. The difference between the two corpora, however, was not resolvable at this sample size (+1 point, 95% CI [-4.0, 5.9] at 14 s; +2 [-0.7, 4.5] at 28 s), indicating that pre-training on behavioral video drives the transfer benefit while corpus diversity contributes no measurable additional gain. Importantly, these accuracy enhancements stem entirely from optimized weight initialization. Because the network architecture remains untouched, feature distillation incurs zero additional computational overhead or latency penalty during online execution, preserving the sub-acquisition inference speeds necessary for real-time experimental control.

## DISCUSSION

Here we present B-REAL, a supervised machine learning platform that achieves both direct video-to-behavior mapping and real-time behavioral classification with end-to-end inference latencies below 15 ms, delivering exceptional performance across both complex social interactions and isolated animal assays. By inserting parameter-free Temporal Shift Module (TSM) blocks directly into an EfficientNet backbone, B-REAL learns spatiotemporal features from raw pixels, with no intermediate keypoint representation, yielding per-frame behavioral predictions on either a GPU or a standard CPU. Because classification keeps pace with acquisition, behavior no longer has to be scored after the session ends: it can trigger environmental changes, drug delivery or neural perturbations while it is still underway.

These properties follow from two architectural choices. EfficientNet backbones were designed by scaling network depth, width and input resolution together, so that each member of the family delivers high accuracy for its computational cost (Tan and Le, 2019); they supply rich spatial features cheaply enough to run within the frame interval even on a CPU, and the b0–b7 family gives users a graded trade between accuracy and speed that can be matched to a rig’s latency budget before any training. Temporal Shift Modules then add temporal processing without adding parameters or computation (Lin et al., 2019): by exchanging a fraction of channels between neighboring frames, every 2D convolution also mixes information across time, so the network models motion as well as appearance at the cost of an image model. Alternatives force a choice between the two — 3D convolutions (Carreira and Zisserman, 2017) and video transformers (Assran et al., 2025) model time explicitly but multiply inference cost, while frame-by-frame image models are fast but see no motion. Consistent with this, lengthening the clip improved accuracy within a backbone (Figure 2D), and B-REAL was the only method to detect fly charges reliably, most of which last a single frame (Figure 4).

Manual frame labeling, rather than model training time, remains the rate-limiting bottleneck in behavioral quantification (Sturman et al., 2020; Luxem et al., 2023). With as little as tens of seconds of annotated video, B-REAL achieves high classification performance exceeding both A-SOiD (Tillmann et al., 2024) and CBAS (Perry et al., 2025). This efficiency is further improved by bypassing keypoint pose estimation training and prediction. Beyond removing manual annotation requirements, keypoint-free classification prevents tracking error propagation and resists common failure modes like physical occlusion or variable illumination conditions (Carreira et al., 2017; Skovorodnikov et al., 2025). While critical, these practical gains understate a more fundamental difference in what information reaches the classifier. A traditional two-stage pipeline reduces the representation of behavior to a set of spatial body landmarks, stripping away all non-skeletal video context before behavioral classification occurs. As a result, actions defined principally by the subject’s interaction with its surrounding environment are poorly captured, because the environment is omitted from the representation. This category of behavior is broad and encompasses core behaviors central to modern neuroscience research. Consummatory actions—such as feeding, drinking, or lick-spout contact—are determined by physical interaction with a specific resource rather than a unique posture, are often brief, and typically unfold within a localized area where spatial context provides essential evidence. Similar constraints apply to nest building, foraging, object investigation, digging, and operant tasks like nose poking or lever pressing. Kinematic trajectories from a feeding mouse and one simply oriented toward a food hopper can be nearly indistinguishable; the pixel-level details are not. Even where keypoint tracking is feasible, as for rigid objects rather than deformable ones like bedding or food, pose-based models require annotating each additional object.

Two insights from our distillation findings deserve emphasis for future deployment. First, feature extraction from the teacher model requires no human annotations, allowing the pre-training corpus to be assembled flexibly from arbitrary, unlabeled recordings. Second, these feature improvements generalized to an experimental perspective entirely absent from the pre-training data: a ventral, bottom-up view, when every pre-training source was an overhead or side-view recording. However, a more heterogeneous pre-training corpus did not yield a measurable advantage over CalMS21 video alone: on the ventral view the two were indistinguishable, and on CalMS21 itself the broader corpus gave smaller gains that never cleared zero. These findings suggest that pre-training on behavioral video is the primary driver of feature transfer, while the value of broad dataset diversity remains to be established.

Sub-acquisition inference latency transforms behavior from an observational output into an active experimental variable. Closed-loop reinforcement of a spontaneous, unconstrained movement like grooming requires detecting it close to its onset; rewarding grooming as B-REAL detected it raised its rate across sessions, showing that the classifier’s output was fast and specific enough to serve as a reinforcement signal. More broadly, any user-defined behavior can now serve as a real-time trigger signal, bypassing the traditional reliance on physical hardware proxies (e.g., beam breaks, microswitches, or lickometers), and going beyond pose-based closed-loop systems, which trigger on a body configuration rather than a classified behavior (Kane et al., 2020; Schweihoff et al., 2021). The duration criterion that turns a detection into a trigger is itself a free parameter: ours required strictly consecutive positive frames, so a single missed frame reset it, and for brief behaviors a windowed criterion (e.g. 4 of the last 6 frames) may trade some specificity for sensitivity — a rule we have not tested, and whose suitability will depend on the behavior and the classifier’s frame-level reliability. This infrastructure opens new avenues for investigating questions previously difficult to address experimentally: shaping naturalistic action sequences like operant responses, tracking circuit-level neural dynamics as actions become goal-directed, and determining how rapidly subjects detect novel contingencies applied to spontaneous behaviors.

As a caveat, our closed-loop validation was conducted using a single testing rig, one mouse strain, and a limited cohort (n = 4), serving as a functional proof-of-concept rather than an exhaustive parameter exploration. In our hardware setup, water was dispensed the moment the frame criterion was met, so the audible solenoid click followed each detected bout immediately and served as a conditioned reinforcer, as in conventional operant shaping; we did not test how the effect depends on the delay between detection and reinforcement, which will matter for rigs with slower reward delivery or longer duration criteria. Additionally, the evaluation of our largest backbone rests on limited data: performance metrics for the EfficientNet-b7 variant reflect a single training run, and two of its three clip configurations exhibited training instability, leading us to report it as an observational benchmark rather than a recommended operating point. Finally, B-REAL models are trained for specific recording geometries and behavioral targets, and we have not systematically tested cross-rig transfer without re-labeling; our distillation results demonstrate how rapidly a new model can be fine-tuned, not the direct reuse of existing weights across distinct setups.

## ACKNOWLEDGEMENTS

We thank members of the Yttri Lab, Drs. Matthew Geramita, Timothy Murphy, and Tony Fong for assistance reviewing the manuscript. Funding provided by NIH BBQS Grant R34DA059509 and Binational Science Foundation Grant 2021296. B-REAL name provided with help from Cheryl Lynn.

## EXPERIMENTAL PROCEDURES

### Animals

For closed-loop behavioral assays, four adult C57BL/6J mice (n=4; 2 males, 2 females) were used. Mouse health and weight were monitored daily and water intake was controlled to 1-2 ml per day, received either within training sessions or in their home cages on break days. Mice were held in a temperature and humidity controlled room maintained on a reversed 12 hour light/dark cycle. While sex was counterbalanced, no sex-based differences were tested given the small sample size, potentially limiting this study’s generalizability. All experiments and procedures were approved by the Carnegie Mellon University Institutional Animal Care and Use Committee. Publicly available benchmark data (CalMS21, Fly-vs-Fly and the pre-training corpora) were collected and approved independently of this study.

### B-REAL model architecture and training

The network core leverages an EfficientNet backbone (Tan and Le, 2019), pre-trained on ImageNet-1k (Deng et al., 2009) and augmented with Temporal Shift Modules (TSM; Lin et al., 2019) inserted prior to every inverted residual block across stages 1 through 7. Video input clips consisting of T frames with shape (B, T, C, H, W)—corresponding to batch, temporal frame index, color channel, height, and width, respectively—are reshaped into standard two-dimensional spatial representations (B·T, C, H, W) prior to convolution. Note that for real-time application we necessarily use a batch size of one. Each TSM unit shifts a designated fraction of channel values along the temporal axis before spatial convolution. Channels are shifted in both directions: one fraction is copied from the preceding frame and an equal fraction from the following frame, with zeros padded wherever the source frame falls outside the clip—the earliest frame for the forward shift, and the final frame for the backward shift. Because the classifier reads out only the final frame of the clip, and that frame’s backward-shifted channels are zero by construction, the prediction at frame t depends exclusively on frames at or before t. The network is therefore causal at the predicted frame, and training and real-time deployment present the network with identical inputs. Crucially, TSM implementation introduces zero additional learnable parameters or extra convolutional layers.

As described in the main text, the 1280-dimensional vector corresponding to the features at time *t* were extracted. To ensure predictions correspond explicitly to the current frame *t* rather than an aggregated temporal clip, only the final frame vector is retained. Classification logits are produced via a lightweight head comprising a dropout layer (p = 0.2) and a linear projection layer. Layers preceding the initial TSM module, corresponding to the convolutional stem, remain frozen during optimization. We did not perform any hyperparameter tuning in any analysis nor evaluation benchmark.

Models were optimized using AdamW (Loshchilov and Hutter, 2019; learning rate 1e-4, weight decay 1e-3) with a cosine annealing schedule ending at 1e-6, paired with cross-entropy loss weighted by inverse class frequency. Boundary-aware label smoothing (ε = 0.3, 5-frame window) was applied to frames surrounding annotation transitions to account for human observer variability. Input images were resized to the size which corresponded to the given EfficientNet backbone, normalized with ImageNet statistics, and augmented with random rotations (±180°), brightness/contrast shifts (±0.2), color jitter, and Gaussian noise. Model checkpoints were selected based on validation Matthews correlation coefficient (MCC) to ensure robust optimization under severe class imbalance.

### Latency profiling and deployment pipeline

Because inference latency is governed by network structure rather than parameter weights, deployment speed can be profiled prior to model training. Candidate architectures are evaluated on synthetic tensor inputs matching runtime dimensions, exported to ONNX format, and compiled using hardware-optimized runtimes: TensorRT for NVIDIA platforms or OpenVINO for Intel architectures. Profiling latency directly within these target deployment formats is critical, as execution timing differs substantially from native PyTorch. The 95th and 99th percentile inference latencies are benchmarked against the real-time frame budget (16.7 ms for 60 fps; 33.3 ms for 30 fps), selecting the maximum viable backbone architecture for final training. Per-clip network latency was calculated as the mean timing across 1000 iterations over 7 independent process executions. All TensorRT configurations were stable across launches (relative standard deviation ≤ 1.2%); three of nine OpenVINO configurations were not (5.5–29.3%), which re-benchmarking traced to CPU frequency scaling: under the default powersave governor, processes launched before the cores ramp up run at roughly half clock, which affects only the fastest configurations, whose entire launch fits inside the ramp (b0 at a 3-frame clip: 33% across all launches, 1.2% excluding the two warm-up launches). The artifact can only inflate latency, so it is conservative for the real-time claims: the slowest CPU configuration was measured precisely (24.45 ± 0.17 ms), and every affected configuration exceeds 30 FPS at the lower bound of its interval (worst case 50 FPS). Fully trained models were exported to TorchScript for direct execution within Python control scripts or Bonsai dataflow environments (Lopes et al., 2015).

### CalMS21 benchmark evaluation

The CalMS21 dataset (Task 1; Sun et al., 2021) was converted into frame-indexed paired images and one-hot target vectors spanning four behavioral categories: attack, investigation, mounting, and other. Video frames were cropped to a 570×570 square, positioned horizontally to center the bounding box of both mice’s keypoints (the pose estimates provided with the dataset); a square of this size, the full frame height, encloses both subjects in 99% of social interaction frames and keeps the geometry undistorted for 180° rotation augmentation. For the cropping ablation (Figure 2E), whole 1024×570 frames were instead resized directly to the network input, with all other settings unchanged. Pre-cropped image caches were maintained at native 570×570 resolution and resized dynamically during training. Evaluation adhered strictly to the official video-level partition (70 training, 19 test videos); frame-level mixing across splits was prohibited to prevent temporal leakage. Validation was conducted on 10 held-out training videos selected to match global class distributions.

To construct label-efficiency curves, training subsets representing fixed durations of annotated video (30 s, 1 min, 5 min, 10 min, and full training set) were sampled at random across the 70 training videos. B-REAL was evaluated across eight independent label draws per budget and A-SOiD (Tillmann et al., 2024) and CBAS (Perry et al., 2025) across three, scoring balanced accuracy (mean per-class recall) on the official test set. For the distillation comparison, each draw fine-tuned both the ImageNet-initialized and the distilled backbone on the same labelled frames, so the two are compared as paired differences. To rule out frame-level overlap between draws, the sweep was repeated with video-disjoint draws: the 70 training videos were partitioned into seven groups and each draw sampled only from its own group’s videos. Two groups contained too few videos with attack frames for any budget sample to include that class and were excluded, leaving five draws per budget. Dense frame caching enabled arbitrary sparse frame sampling while preserving complete T−1 context sequences. Additionally, EfficientNet backbones (B0, B3, B7) were benchmarked across varied temporal clip lengths (3, 5, 7 frames) under a fixed 20-minute training allocation (36,000 frames) over 8 training epochs with early stopping. Performance was compared directly against A-SOiD (a keypoint-based offline classifier) and CBAS (a keypoint-less model fitting an LSTM over frozen DINOv2 embeddings; Oquab et al., 2024) using identical data splits and training budgets. CBAS was given the same keypoint-cropped frames as B-REAL, written out as videos with identical crop geometry.

#### Fly-vs-Fly benchmark evaluation

The aggression subset of Fly-vs-Fly (Eyjolfsdottir et al., 2014) comprises ten 30-minute videos (54,000 frames at 30 fps, 144×144 pixels) of pairs of male *Drosophila*, in which each fly is annotated frame by frame for lunge, wing threat, charge, hold and tussle; fly-frames carrying none of these form the ‘other’ class. The dataset’s official split was used (videos 1–5 for training, 6–10 for testing). Fly-frames that the annotators marked as unscorable (0.43%) were excluded from training and scoring rather than assigned to ‘other’. Where two behaviors were annotated on the same fly and frame (0.13% of fly-frames), the rarer behavior was kept. Because CBAS classifies whole frames and cannot attribute a call to one animal, and B-REAL here also processes the whole frame, all three methods were trained and scored on a single per-frame target: a frame carried a behavior if either fly performed it, with the rarer behavior taking precedence when the two flies differed (charge, then hold, tussle, lunge and wing threat), and a frame was scorable if at least one fly was.

Label budgets of 30 s and 1, 5 and 10 min were each sampled in eight random draws, and each method was also trained once on the complete training set. Draws were sampled from all training fly-frames pooled across the five training videos, so they were not video-disjoint. Budgets were counted at 60 fly-frames per second of video (30 fps × two flies). Each sampled fly-frame contributes its video frame, labelled from both flies’ annotations, and the two flies of a frame were rarely both drawn, so a draw held nearly as many distinct frames as fly-frames (1,790–1,799 at 30 s, 34,754–34,818 at 10 min). A Fly-vs-Fly budget therefore contains about twice as many labelled frames as the same nominal budget on CalMS21. The first two frames of each video, which lack a complete 3-frame clip, were never drawn. A purely random 30-s draw contains on average fewer than one charge frame;. each draw was therefore topped up to at least one example of every behavior, placed in the portion of its video used for training rather than validation, with each addition replacing a randomly chosen ‘other’ frame so that the budget size was unchanged. At the smallest budgets, the single charge example in a draw is present by construction rather than by chance. All three methods were trained on the same frames in every draw.

B-REAL used the configuration of Figure 2B and C (EfficientNet-b0, 3-frame clip, ImageNet initialization), with the trainer and its model and data-loading settings copied unchanged from CalMS21. Frames were decoded to greyscale and left uncropped at their native 144×144: the videos are already cropped to the chamber, and a tighter crop around each fly would discard the two flies’ relative position, which defines behaviors such as lunge and wing threat. Training ran for up to 100 epochs, with early stopping after 8 epochs without improvement in validation MCC and checkpoint selection by the same measure; the validation set was the chronologically final 15% of each training video’s drawn frames.

A-SOiD was given five keypoints per fly derived from the dataset’s ellipse-and-wing tracks: the body centroid, the head and tail at either end of the body’s major axis, and the two wing tips, ten keypoints in all; tracking gaps (0.03% of fly-frames) were filled by linear interpolation. A-SOiD bins its pose features in time, with the bin length set by the shortest behavior of interest. Because more than half of charge bouts last a single frame, this was set to one frame (minimum duration 0.05 s); the default 0.1 s gives 3-frame bins and removes 39% of charge frames before training. At this setting A-SOiD applies no temporal smoothing to its features. A-SOiD’s random-forest hyperparameters were used unchanged, and its active-learning loop was run within each draw, so it could select only from labels the draw contained: the forest was seeded with 10% of each class from 80% of the draw (the remainder is held out by A-SOiD for scoring its iterations), and up to 20 of the least confident non-’other’ examples (maximum class probability below 0.5) were added per iteration for up to 200 iterations. The seed fraction was raised from A-SOiD’s default of 1%, which at 30 s seeds about 13 examples, all classified confidently, so that the loop stops at once and predicts only ‘other’; it was held at 10% for every draw so that only the budget varied. A-SOiD’s final forest was therefore trained on 8–10% of the frames in each draw. CBAS was trained on the raw videos, encoded with its default DINOv2 encoder (with registers, base size), using the training defaults of its graphical interface (31-frame windows, a single-layer LSTM with 64 hidden units, up to 10 epochs, and the best of 3 × 5 runs selected by weighted F1 on a 15% validation split of the draw).

All methods were scored on the per-frame target of the five test videos: 269,902 frames for B-REAL, which excludes the first two frames of each video, and 269,762 for CBAS, which loses a few frames at the edges of each 31-frame window. A-SOiD’s one-frame bins map directly onto frames; its scoring also counted the 88 test frames on which neither fly was scorable, as ‘other’ (0.03% of test frames). Balanced accuracy was computed as the mean recall over all six classes, including ‘other’, and per-class F1, precision and recall were reported alongside it.

#### Bounding box subject cropping

To isolate moving subjects, an optional spatial cropping routine was implemented using background subtraction rather than deep learning-based detection. A reference background model was generated by averaging *k* subject-free initial frames, most commonly 500. Incoming video frames were converted to greyscale, downsampled, and subtracted from the background reference. In directional mode, only pixels darker than the background were preserved to isolate dark subjects on light surfaces, whereas absolute mode captured pixel intensity changes in either direction to accommodate arbitrary coat colors. Target difference masks were thresholded, cleaned via morphological operations, and fitted with a fixed-size square bounding box centered at the primary contour centroid. Centroid trajectories were smoothed using an exponential moving average with a dead-band threshold to suppress bounding box jitter, and small artifacts below a minimal area threshold were discarded. This optional module provides lightweight pre-processing, though performance degrades if ambient lighting fluctuates significantly across extended recording sessions.

#### Behavioral testing apparatus and closed-loop platform

Subjects were recorded from below in an open, 12-inch square arena using a single free-running USB camera operating at 1280×720 resolution. Sessions lasted 45 minutes. A fixed spatial region of interest (ROI) spanning 690×685 pixels was assigned to encompass the arena, alongside manually calibrated white balance, exposure, and analog gain settings. Online behavioral processing and classification were bound to a 30 FPS execution limit, which was consistently maintained across all experimental sessions and subjects.

Grooming (head and face, but not body) was categorized using a dedicated binary B-REAL model (B0 backbone, 3-frame context) trained on manually annotated video sequences. At every image frame, the B-REAL classifier computed a pseudo-probability indicating the likelihood the animal was currently performing the behavior of interest. An internal frame counter incremented sequentially upon each positive classification and immediately reverted to zero upon encountering any single negative frame. When the consecutive frame counter satisfied the required temporal threshold—defined as the ceiling of the target duration divided by the 33.3 ms frame period—a single trigger byte was transmitted to the microcontroller. Consequently, reward delivery required uninterrupted, continuous execution of the target behavior rather than a moving-window majority vote.

A reward port equipped with an infrared photobeam sensor was permanently fixed through the arena wall. Visual feedback denoting reward availability was delivered via an overhead white light positioned directly above the port, as well as additional auditory feedback from the activation of the solenoid. Control signals were transmitted to the system microcontroller via a dedicated serial communication link configured at 9600 baud.

The system hardware operated across three distinct functional regimes. In the initial armed configuration, receipt of a trigger signal prompted immediate fluid delivery via a brief 50 ms solenoid actuation, dispensing the reinforcement prior to port arrival. Following solenoid closure, the overhead flood light illuminated as the apparatus transitioned into a locked configuration, suppressing subsequent trigger inputs to ensure that concurrent behavioral criteria generated no additional reward or visual signaling while a delivery remained uncollected. Photobeam disruption at the reward interface extinguished the visual signal and restored the armed state for subsequent trial sequences. Consequently, reinforcement depended exclusively on execution of the target action rather than approach kinematics; active illumination indicated dispensed reward availability rather than arrival-gated delivery. Individual light cycle durations thus mapped directly to single earned rewards, providing the precise interval for calculating retrieval latencies. Temporal criterion thresholds were maintained at 100 ms (corresponding to 3 consecutive frames) throughout grooming blocks and early rearing paradigms, subsequently expanding to 500, 750, or 1000 ms (reaching 1500 ms in terminal experimental stages) across late-stage rearing protocols.

#### Data logging and telemetry

System metadata—including subject identifier, behavioral target, active network model, operational mode, duration threshold, and epoch durations—was logged at session initiation. Sequential cue-state transitions were logged frame-by-frame, recording state 1 upon meeting the consecutive-frame criterion and state 0 on the subsequent negative frame. The resulting session log recorded trigger events and post-criterion extensions rather than raw frame-by-frame behavioral states; consequently, all global behavioral quantification was performed post hoc using offline network scoring, reserving session logs strictly for trigger latency analyses

#### Foundation model representation distillation

To enhance temporal feature representations beyond static ImageNet initializations, representations were distilled into the B-REAL backbone from a frozen V-JEPA 2 ViT-L video foundation model (Assran et al., 2025) operating on 16-frame inputs. Representation quality was assessed by linear probing: each backbone was frozen, a multinomial logistic probe was fitted on 40 videos disjoint from the test set, and balanced accuracy was scored on CalMS21’s official 19 held-out videos. A single anchor set was drawn once and shared by every candidate, so all comparisons are paired. Balanced accuracy rose from 0.623 for the ImageNet-initialized backbone to between 0.710 and 0.754 across the twelve distillation epochs. This probe was used only to characterize the representations and to validate the checkpoint selectors described below; it never itself selected a checkpoint, as fitting a probe on held-out video would reintroduce the selection bias LogME (You et al., 2021) is used to avoid. Cross-model feature regression confirmed that less than half of the teacher’s variance (R² = 0.47) could be captured by ImageNet features, demonstrating substantial novelty in the target representations. Teacher feature maps were pre-computed and stored on disk, framing the student distillation task as a supervised feature regression problem without risk of representation collapse.

Pre-training utilized an unlabelled, multi-source mouse video corpus spanning five diverse experimental settings (JAX (Geuther et al., 2019), CalMS21 training set, CRIM13 (Burgos-Artizzu et al., 2012), lockbox dataset (Reiske et al., 2025), and MABe22 (Sun et al., 2023)) covering varied camera angles, arena geometries, illumination conditions, and coat colors. All source videos were standardized to 30 fps. Multi-animal bounding boxes were constructed dynamically around the union of detected subject contours using background subtraction without utilizing behavioral annotations. Crucially, the 19 CalMS21 test videos were fully excluded from pre-training. A second student was distilled from unlabelled CalMS21 training video alone, as a control for corpus diversity. The CalMS21 budget sweep compared the two directly; the ventral-view transfer used a CalMS21-only student subsampled to the same number of anchors as the five-source corpus (27,579 and 27,214), so that the two differ in diversity but not in quantity. Student network clips (frames *t*−2, *t*-1, and *t* at 224 px) were aligned to end on the final frame of the teacher window (frames *t*-15 through *t* at 256 px). Pre-trained checkpoints were selected via LogME scoring on training labels and fine-tuned on target datasets using paired multi-seed comparisons against standard ImageNet-initialized baselines.

#### Ventral-view transfer benchmark

Transfer to a recording geometry absent from pre-training (Figure 5C, D) was tested on a single annotated video of one mouse filmed from below; no part of this video was included in any pre-training corpus. Frames were cropped to a 220×220 pixel colour window centred on the animal by the background-subtraction routine described above. Each of the first 35,278 frames (19.6 min at 30 fps) was annotated with one of eight classes: other, locomotion, right turn, left turn, body groom, face/head/paw groom, rear and itch. Turn direction refers to the animal’s own left and right. Frames labelled rear or left turn were removed rather than reassigned, leaving six classes and 29,589 frames, with class prevalence ranging from 3.2% (itch) to 57.5% (other).

The video was divided chronologically, without shuffling, into training (first 70%), validation (next 15%) and test (final 15%; 4,437 frames) blocks. Frames whose 3-frame clip would extend into the preceding block were excluded, so no clip drew context from across a split boundary. Labelled-data budgets were 2% and 4% of the training block (413 and 829 frames, or 14 and 28 s at 30 fps), sampled at random separately within each class so that every subset kept the class proportions of the full training block. Each budget was sampled in eight independent draws, and each draw fine-tuned all three initializations (ImageNet, the CalMS21-only student and the five-source student) on the same frames, so they are compared as paired differences. At each budget, two of the eight draws had been trained twice on identical frames; the two runs were averaged so that each draw contributes one value. The validation and test blocks were not subsampled. Fine-tuning used the CalMS21 configuration (b0 backbone, 3-frame clip, and the optimizer, augmentation and boundary-aware label smoothing described above) for up to 100 epochs, with early stopping after 15 epochs without improvement and checkpoint selection by validation MCC; balanced accuracy was scored on the test block. Both students were taken from the same distillation epoch, fixed in advance rather than selected by LogME, so neither the transfer video nor its labels influenced the choice of initialization.

#### Statistical analysis

All tests are two-sided. Where tests are corrected for multiple comparisons, we used the Holm–Bonferroni procedure: comparisons of B-REAL against the baseline methods are corrected over the two baselines compared at each budget and metric, the distillation comparisons marked in Figure 5B and 5D over all tests in each panel (eight in 5B, six in 5D), and the bout-duration comparison in Figure 4F–H over the four duration bands tested. Confidence intervals are unadjusted. All other p-values are uncorrected. Classifier performance (Figure 4B, C) is reported as mean ± s.d. over three training seeds; each seed’s detection delay is its median onset-to-detection delay. In the closed-loop experiment, each animal’s grooming rate was expressed as percent change from its own mean over grooming-block sessions 1–3, and acquisition was tested with a one-sample t-test of each animal’s mean change over sessions 7–9 (the last three shown in Figure 4D; every animal ran at least ten) against zero (n = 4 animals), which equals a paired t-test of sessions 7–9 against sessions 1–3 on this percent-change scale. Sessions 1–3 were already rewarded, so the change is measured from a rewarded starting point rather than a pre-reward baseline. At session resolution, each session’s rate was centred on its own animal’s mean, and the early and late halves of each animal’s grooming block (the middle session of an odd-length block counted as late) were compared with a two-sample Student’s t-test (n = 54 sessions), in place of a mixed model with a per-animal intercept. For criterion bout durations (Figure 4F–H), sessions 1–3, 4–6 and 7–9 were pooled within each animal; the share of bouts in each of four duration bands (100–133, 167–300, 333–500 and >500 ms, chosen by visual inspection) was compared between sessions 7–9 and 1–3 with a paired t-test over animals (n = 4). Bands in Figure 4D are 95% t-intervals across animals; bands in Figure 4F and 4G are s.e.m. across animals. On CalMS21, B-REAL (eight label draws per budget) was compared with each baseline (three seeds) by Welch’s t-test at each budget. On Fly-vs-Fly, all three methods were trained on the same eight label draws per budget, and paired differences were tested with the Wilcoxon signed-rank test (with eight draws, the smallest attainable value is p = 0.008 before correction and 0.016 after, for both baselines under Holm). For distillation, each draw fine-tuned the ImageNet-initialized and the distilled backbone on the same labelled frames; per-draw paired differences are reported as mean with 95% t-interval and tested against zero with a one-sample t-test, Holm-corrected as above.

#### AI Usage

Claude Opus 5 (Anthropic, 2026) was used for code generation and evaluation as well as refinement of the manuscript text. The authors assume all responsibility for results and the final work.

## CODE AVAILABILITY

Code is available publicly at https://github.com/adeneagle/B-REAL.

