## Supplemental figures for "B-REAL: A keypoint-less behavior platform for offline classification and real-time detection"

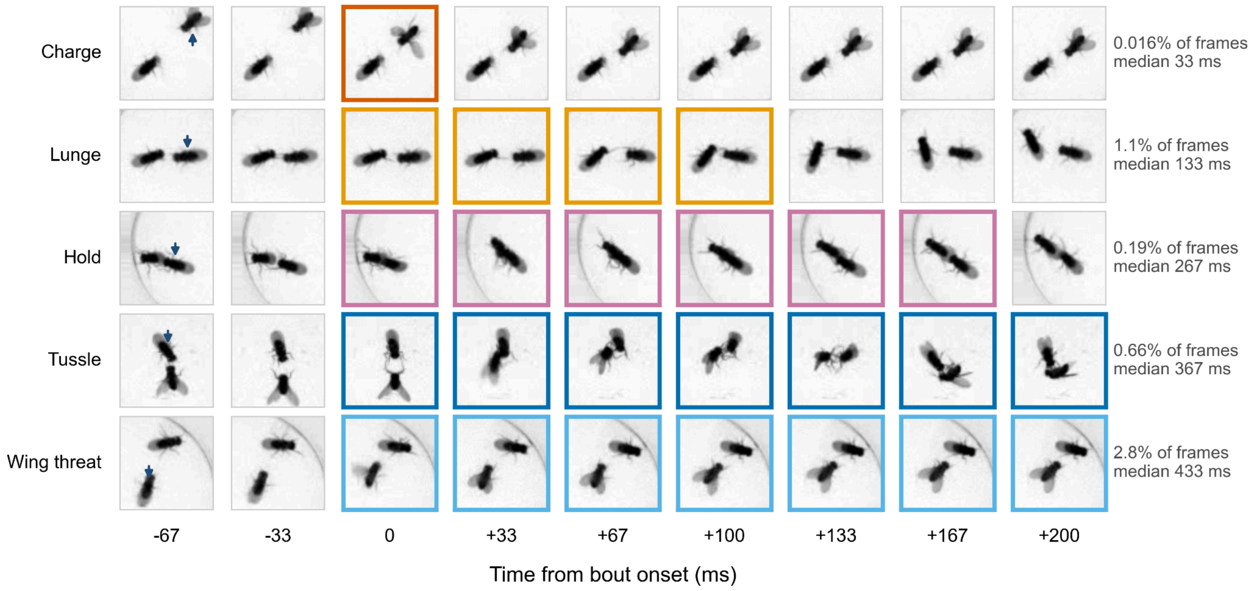


**Supplementary Figure S1. All five Fly-vs-Fly aggression behaviors.**
Nine consecutive frames (30 fps, 33 ms apart) around one annotated bout of each behavior, cropped around the pair and aligned so that the bout begins at 0 ms. Outlined frames are those manually annotated with the given behavior, and the arrowhead in the first frame marks the fly to which the label belongs. Right, each behavior's share of all scored fly-frames and its median bout duration over all ten movies. Charge occupies a single frame (0.016% of frames, median 33 ms) and lunge four (1.1%, 133 ms), while hold (0.19%, 267 ms), tussle (0.66%, 367 ms) and wing threat (2.8%, 433 ms) last several times longer. Examples were chosen by eye from typical-length bouts for legibility and are illustrations. Contrast scaled separately for each row.

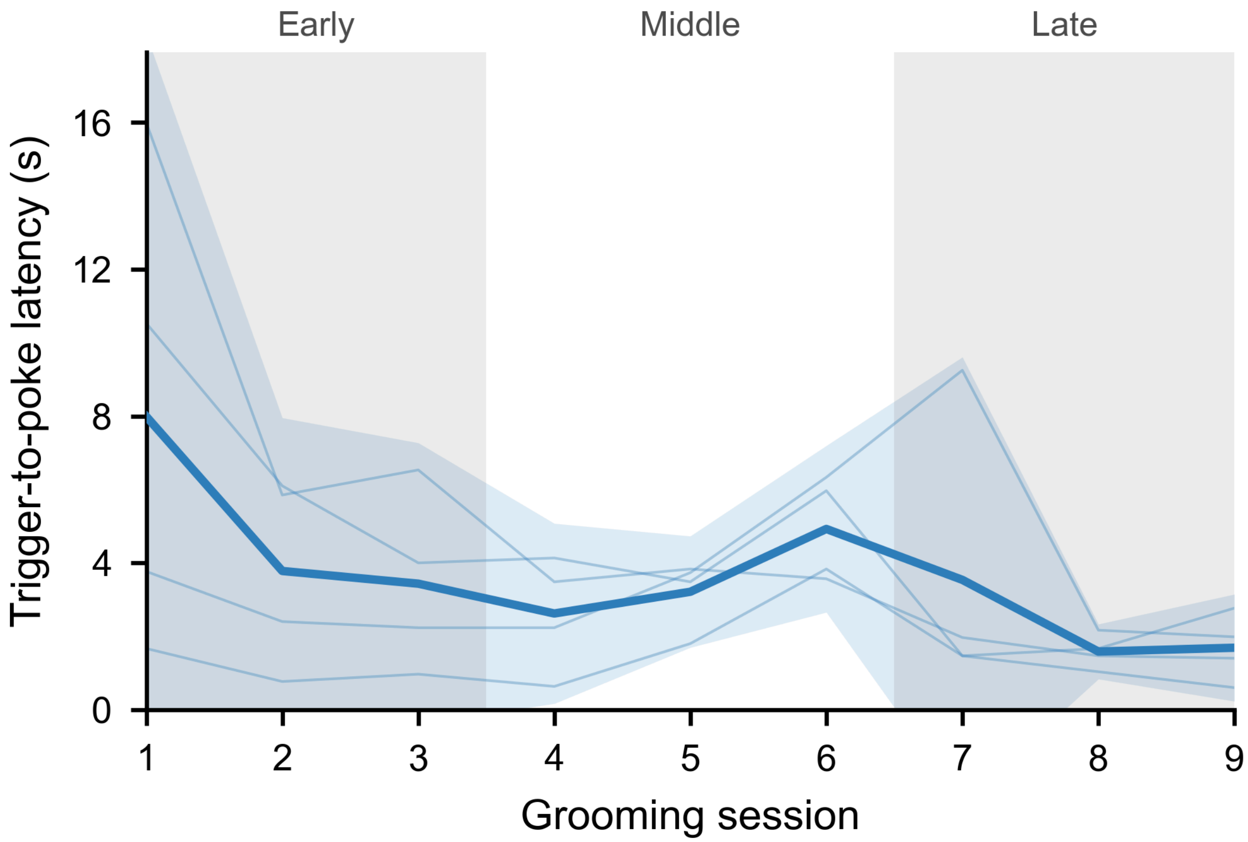


**Supplementary Figure S2. Reward receipt latency decreases over training.**
Median latency from reward delivery to the next nose-poke at the reward port, in each of the first nine grooming sessions (n = 4 mice; the sessions of Fig. 4d). Water was delivered when the online classifier detected grooming on three consecutive frames; detections that arrived while an earlier reward was still uncollected delivered no water and were not counted. Nose-pokes were detected from the session video. Thin lines, individual animals; thick line, mean; band, 95% CI across animals; grey shading, the early (sessions 1–3) and late (7–9) windows of Fig. 4d. Latency fell from session 1 to session 3 in all four animals, to 0.48 of its session-1 value (95% CI 0.33–0.70; paired *t*-test on log latency, $p = 0.008$). Reward delivery was audible via a ‘click’ from the solenoid.
